# Temporally integrated transcriptome analysis reveals ASFV pathology and host response dynamics

**DOI:** 10.1101/2022.05.06.490987

**Authors:** Lin Lv, Tianyun Zhang, Yanyan Zhang, Asif Ahsan, Xiaoyang Zhao, Zhiqiang Shen, Teng Chen, Ning Shen

**Affiliations:** Department of Hepatobiliary and Pancreatic Surgery, First Affiliated Hospital, College of Medicine, Zhejiang University, Hangzhou, Zhejiang Province, 311121, China; Liangzhu Laboratory, Zhejiang University Medical Center, Zhejiang University, Hangzhou, Zhejiang Province, 311121, China; Changchun Veterinary Research Institute, Chinese Academy of Agricultural Sciences, Changchun, Jilin Province, 130122, China; Shandong Binzhou Academy of Animal Science and Veterinary Medicine, Shandong Academy of Agricultural Sciences, Binzhou, Shandong Province, 256600, China; Shandong Lvdu Bio-Sciences and Technology CO., LTD, Binzhou, Shandong Province, 256600, China

## Abstract

African swine fever virus (ASFV) causes a lethal swine hemorrhagic disease and is currently responsible for widespread damage to the pig industry. The molecular mechanisms of ASFV pathogenicity and its interaction with host responses remain poorly understood. In this study, we profiled the temporal viral and host transcriptomes in porcine alveolar macrophages (PAMs) infected at 6, 12, 24 and 48 hours with highly virulent (SY18) and low virulent (HuB20) ASFV strains. We first identified profound differences in the virus expression programs between SY18 and HuB20, while the transcriptome dynamics in host cells were dominated by infection time. Through integrated computational analysis and experimental validation, we identified differentially expressed genes and related biological processes, and elaborated differential usage of the NF-kappaB related pathways by the two virus strains. In addition, we observed that compared to the highly virulent SY18 strain, HuB20 infection quickly activates expression of receptors, sensors, regulators, as well as downstream effectors, including cGAS, STAT1/2, IRF9, MX1/2, suggesting rapid induction of a strong immune response. Lastly, we constructed a host-virus coexpression network, which shed light on pathogenic functions of several ASFV genes. Taken together, these results will provide a basis for further mechanistic studies on the functions of both viral and cellular genes that are involved in different responses.

**Author Summary:** Since it was first described in Kenya in 1921, ASF has spread across sub-Saharan Africa, the Caribbean, the Western Europe, the Trans-Caucasus region, and the Russian Federation. Recent outbreaks have also been reported in Asia, which has devastated the pig industry, resulting in an approximately 40% reduction in pork worldwide. In the absence of effective vaccine or treatment, the mortality for infections with highly virulent strains approaches 100%, while low virulent strains causing less mortality spreads fast recently. Nevertheless, the mechanisms of ASFV pathogenicity, especially the differences between highly and low virulent strains remain poorly understood. Here, we used RNA-seq to analyze the viral and host transcriptome changes in PAMs infected with a virulent strain (SY18) or an attenuated strain (HuB20) at different stages. We found that the presence of ASFV significantly affected the cellular transcriptome profile. In addition, we did temporal and described the dynamic expression programs induced in the host cells by ASFV infection of different virulence strains. In particular, we identified differential gene expression patterns in host innate immune responses and expressed cytokines and chemokines between ASFV strains of different virulence. Our study provides new insights into ASFV pathogenicity research and novel drug or vaccine targets.

## Introduction

African swine fever, caused by African swine fever virus (ASFV), is a fatal hemorrhagic disease of domestic and wild pigs (1-3). Outbreaks of ASF have spread rapidly throughout Eastern Europe, Africa and Asia, making ASF a major threat to the pig industry worldwide, especially in the last decade (4, 5). ASFV is one of the most complex DNA viruses known to date, encoding over 150 proteins involved in a variety of stages of ASFV life cycle, including evasion of host immune response, entry into host cells, RNA modification, DNA repair, and virion assembly (6). Macrophages and monocytes are the primary targets of ASFV and are thought to be critical for virus replication and dissemination (6, 7). Despite extensive research on ASFV and its devastating effects on the host, no effective drug or vaccine is available (4). A major restriction in the development of effective ASFV antiviral therapies is due to the limited understanding of the molecular mechanisms of ASFV transcription and its interaction dynamics with the host cell, i.e., studies of a single gene or pathway of ASFV infection fail to provide a satisfactory understanding of the host-virus interaction dynamics (4, 8, 9). Consequently, comprehensive profiling of ASFV gene expression and its interaction with the host transcriptome is highly valuable, as it may provide novel insights for the development of antiviral therapies and effective vaccines.

RNA sequencing (RNA-Seq) is a high-throughput experiment that can be applied to profile the transcriptome of host and virus during infection (10-13). Using RNA-seq, researchers quantified gene expression levels in Vero cells infected with ASFV-BA71V at early (5 hour) and late (16 hour) stages, providing insights into the temporal expression of known and novel viral genes (14). However, the use of non-ASFV targeted cells is suboptimal and may introduce bias. Another study profiled ASFV transcripts in the blood of pigs infected with either virulent (Georgia 2007, GRG) or attenuated (OURT33) strains. This analysis showed unique gene expression patterns between GRG and OURT33, including host genes associated with macrophages and natural killer (NK) cells, and viral genes associated with modification of host immunity (15). However, a limitation of this study is that they used mixed cell types, thus transcriptomic changes in the host cells may be complicated by secondary effects in uninfected cells. Some studies also applied RNA-seq to describe the gene expression of porcine alveolar macrophages (PAMs) infected with the highly virulent ASFV strain Malawi LIL20/1, Georgia 2007 or CN/GS/2018, where changes in some important cytokines and transcription factors in host cells after ASFV infection were reported (16-18). However, the dynamic transcriptome changes in host cells after ASFV infection, especially the more common low virulent ASFV strains, remain unclear.

Previous studies have demonstrated that inhibition of interferons (IFNs) is a crucial strategy utilized by ASFV to evade immune responses (19-22). The highly virulent ASFV strains can suppress the immune response by encoding genes such as the multigene family 360 (MGF360) and multigene family 505 (MGF505), while attenuated strains, on the contrary, are less studied in this regard (23-26). In particular, differences in the host cell immune response following infection with ASFV of different virulence remain poorly understood. Thus, elucidation of the host immune response of different strains could provide insightful perspectives on ASFV immune evasion strategies and shed light on new vaccine development strategies.

Cytokines and chemokines are critical to macrophage function, such as regulating effective immune responses, and linking innate and adaptive immunity (27-31). As a result, ASFV is known to antagonize immune and inflammatory responses by controlling host cell cytokines and chemokines expression (21). *In vitro* studies on macrophages showed that the low virulent ASFV NH/P68 strain induced high expression of IFN-α, IL-6, TNF-α and IL-12 compared to the highly virulent ASFV L60 strain, while another studies showed that both the NH/P68 and 22653/14 (highly virulent) strains negatively regulated IL-6, IL-12 and TNF-α release in macrophages (32, 33). The conflicting results may be due to differences in the virulence of the strains tested, the dose and duration of infection or sampling timepoints. Therefore, an experimental design to better understand the pattern of cytokines and chemokines changes after ASFV infection will be beneficial to the understanding of ASFV pathogenicity.

In this study, we performed RNA-Seq experiments on PAMs infected with virulent (SY18) or attenuated (HuB20) ASFV strains across multiple stages of virus infection, and profiled the transcriptome of the virus and host respectively. Both SY18 and HuB20 strains belong to type II ASFV, where SY18 strain was firstly obtained from specimens in the initial ASF outbreak in northeastern China and caused almost 100% mortality in pigs. The HuB20 strain is a naturally attenuated ASFV isolated from southern China that causes 30-40% death in infected pigs and is always mild in clinical symptoms. We characterized the temporal expression dynamics of both host and virus genes and enriched functions. In particular, we identified distinct differentially expressed host genes involved in NF-kappaB pathways, differences in the host innate immune response, and distinct expression patterns of cytokines and chemokines in response to ASFV infection between strains of different virulence. Our results help provides insights into a comprehensive understanding of the host-virus interaction dynamics after ASFV infection, as well as the differential expression programs between the virulent and attenuated strains.

## Results

### Landscape of host-virus transcriptome dynamics in two ASFV strains

To study the dynamics of the host-virus transcriptome during ASFV infection, we infected PAMs with SY18 and HuB20, two ASFV strains of different virulence, and profiled their transcriptome through RNA-Seq at 6, 12, 24 and 48 hours post infection (hpi), respectively (Fig 1A). Principal component analysis (PCA) of the virus transcriptome suggests that expressional variation between the two virus strains (strain-specific) dominates the transcriptome variation among all samples, explaining 88% of all variation on PC1, whereas time-course expression variation within virus strains only accounts for 3% variation on PC2. This indicates that expressional differences between the virus strains might play a major role in the virulence of the two virus strains (Fig 1B). In contrast, transcriptome profiling of the host genome identifies time-course changes as a dominant variation, in contrast to strain specific differences, where 70% of the variance aligned with infection timecourse (Fig 1C). This indicates that transcriptional responses on the host cells are not distinctly different between the two virus strains, despite distinct clinical outcomes. Nevertheless, we still observed larger variation in the host transcriptome between two virus strains infected with the same timepoint than between biological replicates of the same condition, suggesting that a differential host response in expression exists with infection of the two ASFV strains.

**Fig 1.**
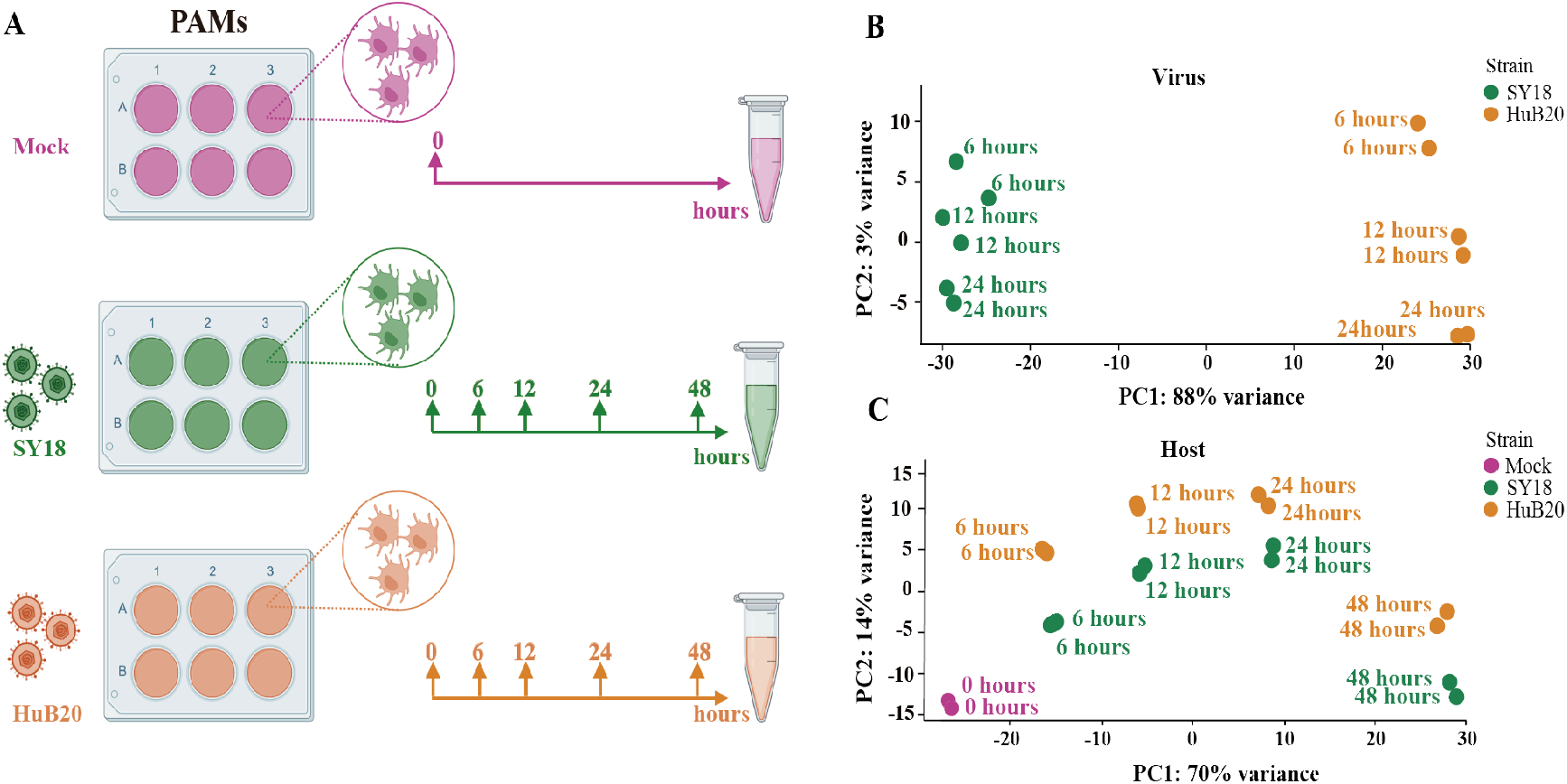
RNA-seq analysis of ASFV strains-infected PAMs. (A) The workflow represents the process of sample collection in this study. PAMs were mock-infected or infected with ASFV strain SY18 or HuB20 (MOI= 3), followed by sample collection at 6, 12, 24 and 48 hpi. Total RNA was extracted and polyA enriched RNA sequencing was performed. The principal component of each sample was analyzed considering the virus genes (B) or host genes (C) expression in the corresponding sample. Samples corresponding to each experimental group were plotted on the first two principal components.

### Viral gene expression programs and functional annotation

To study the viral expression programs of SY18 and HuB20, we plotted the temporal gene expression profiles of all viral genes in a replication cycle (6, 12 and 24 hpi), as indicated by the PCA plot of the two virus strain transcriptomes (Fig S1A and B). Figure 2A demonstrates the viral gene expression profiling of SY18 and HuB20 strains. We identified six clusters according to their expression pattern (S1 Table). Cluster Ⅲ and Ⅵ presented similar expression patterns, while cluster Ⅰ, Ⅱ, Ⅳ and Ⅴ showed distinct expression programs between the two virus strains. Consistent with previous reports involving point mutations and deletions of many genes in HuB20 compared to SY18 (34), our results demonstrated that the dynamic viral gene expression programs between the two virus strains were dramatically different as well.

**Fig 2.**
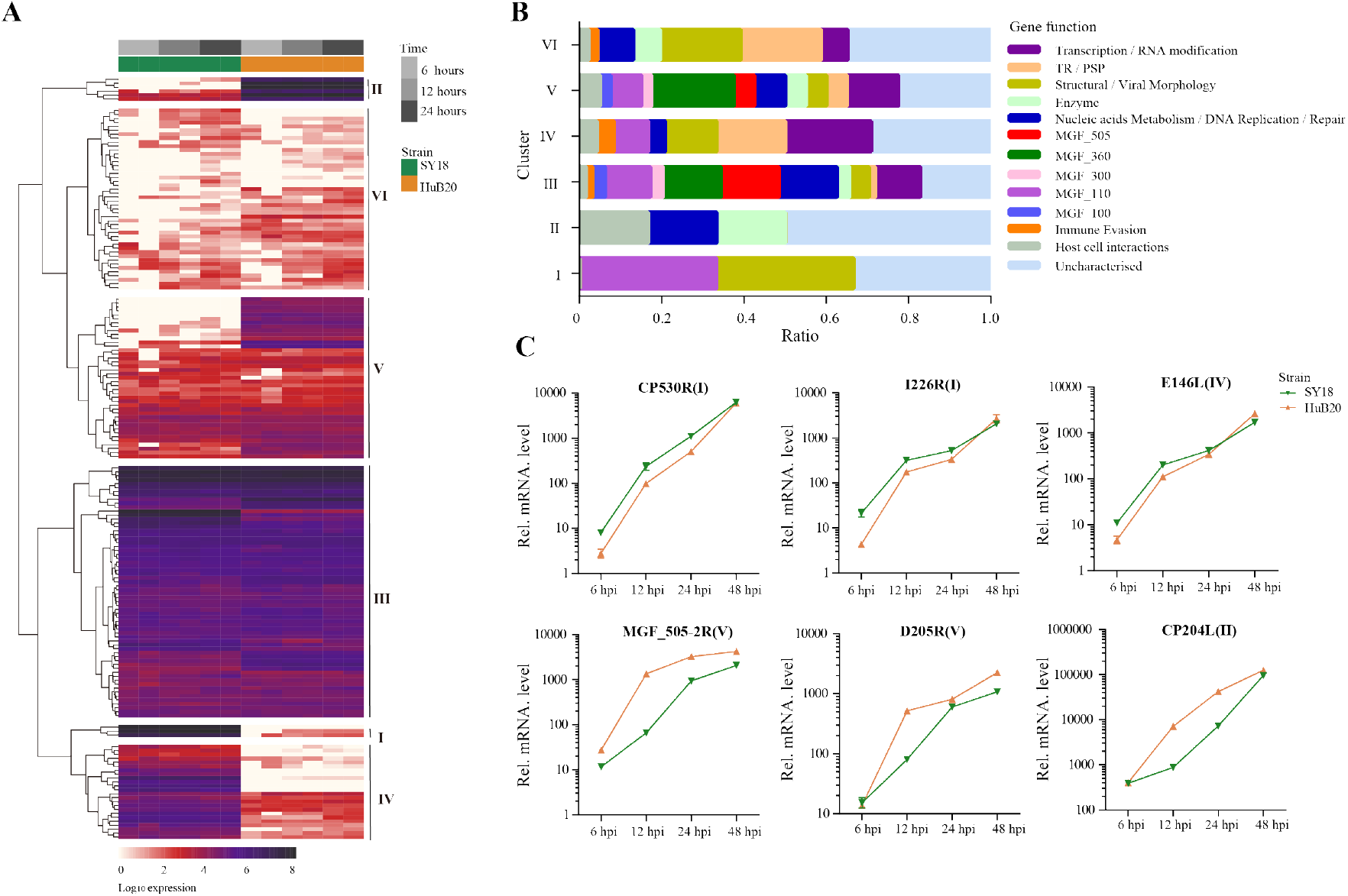
Expression analysis and functional classification of ASFV genes. (A) Heatmap shows the expression levels for the 184 viral genes in the ASFV SY18 and HuB20 strains. (B) The functional classification of the detected 184 ASFV genes in SY18 and HuB20 strains, annotated with the most enriched function and divided into 7 clusters. (C) Validation of randomly selected ASFV gene expression by real-time PCR. At 6, 12, 24, and 48 hours after PAMs were infected with ASFV SY18 and HuB20 strain (MOI= 3), the transcriptional level of CP530R, I226R, E146L (highly expressed in the SY18 strain infected group) and MGF_202-R, D205R, CP204L (highly expressed in the HuB20 strain infected group) were detected by RT-qPCR. The fold-difference was measured by the 2^-ΔΔCt^ method. The RNA levels were normalized to the corresponding β-actin.

Next, we annotated the 184 viral genes with functional groups and profiled the functional composition of different clusters of viral gene expression programs (Figure 2B). Interestingly, cluster III, which contains constitutively highly expressed genes in both virus strains, presented the most versatile functions covering all categories of functional groups. On the contrary, functional groups for genes in cluster VI mainly involves structural/viral morphology, transmembrane region/putative signal peptide (TR/PSP), and DNA replication, with no MGF family genes involved, suggesting that genes associated with viral particle packaging, maturation and propagation were consistently expressed at relatively late timepoints after virus infection for both SY18 and HuB20.

The rest of the clusters encode cluster I and IV, which were expressed higher in SY18, and cluster II and V that were expressed higher in HuB20, respectively. Functional annotations of both sides of gene clusters encompass diverse functional categories, with SY18 high expression cluster IV containing several genes involved in immune evasion, while HuB20 high expression clusters cover more diverse MGF family genes. In addition, we confirmed the differential expression levels of selected viral genes from both sides of the clusters using RT-qPCR (Figure 2C). Thus, our results suggested distinct differences in viral gene transcription programs between the two strains over time. The dynamic viral gene expression programs and functional annotations in our analysis might contribute to the understanding of the cooperative viral gene functions and pathogenicity differences of ASFV strains with different virulence.

### Host transcriptome dynamics after infection of two ASFV strains

In addition to the virus transcriptome, we also analyzed the host transcriptome along different infection stages of the two virus strains. A total of 2320 significant differentially expressed genes (DEGs) (1345 upregulated and 975 downregulated) and 2304 DEGs were identified in SY18 and HuB20 (1321upregulated and 983 downregulated) respectively, compared to mock-infected samples (P < 10^−10^) (Fig 3A). Meanwhile, approximately two thirds of the DEGs were (988 in the upregulated group and 698 in the down-regulated group) shared by the two strains (Fig 3B), confirming that similar level of transcriptome response was stimulated by the two virus strains of different virulence. However, about one third of the DEGs demonstrated specificity between SY18 and HuB20 strains in both upregulated and downregulated group, suggesting that potentially diverse expression programs were involved in the host transcriptome after infection with the two virus strains.

**Fig 3.**
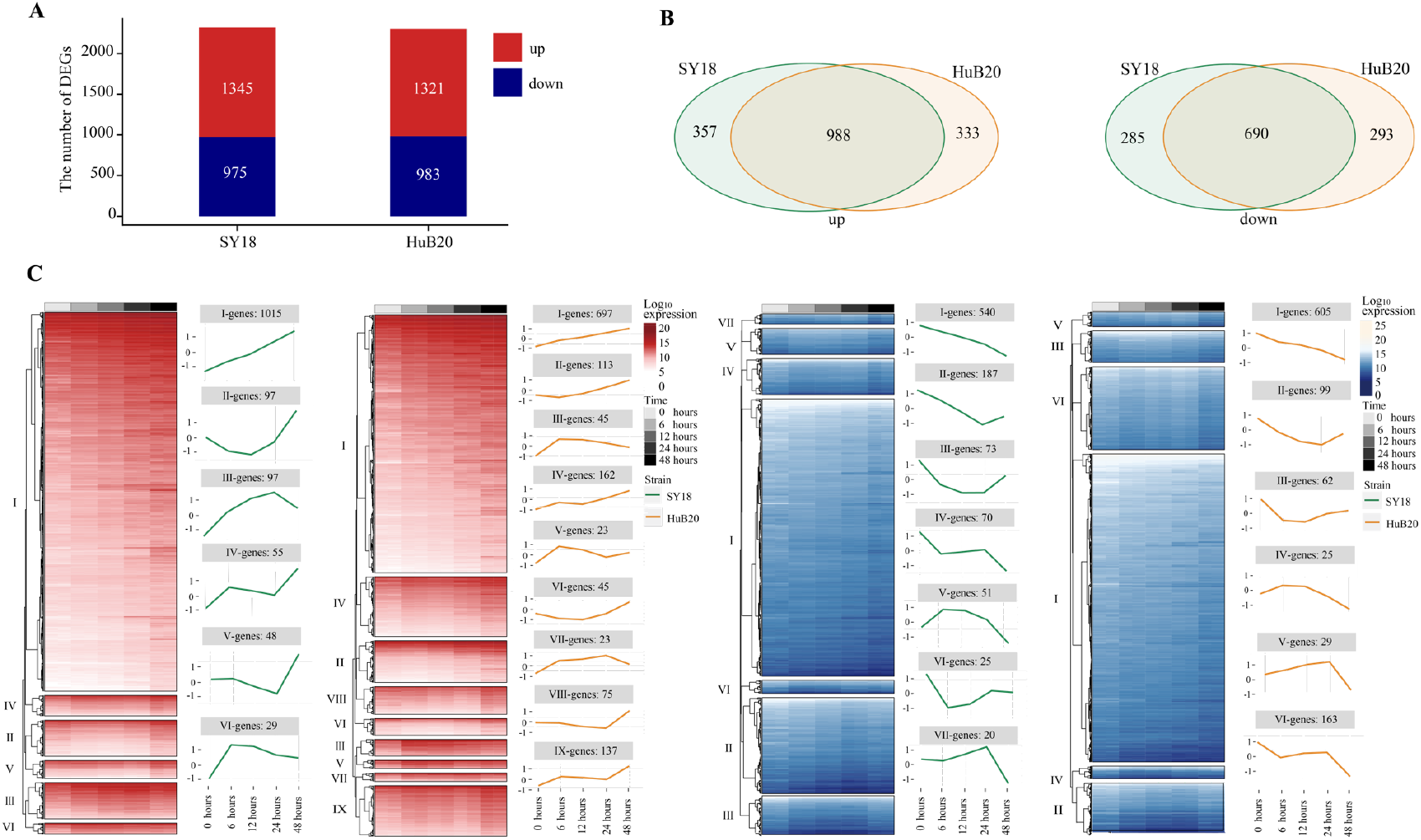
Differentially expressed genes (DEGs) analysis in host with time series. DEGs are examined by the Likelihood Ratio Test (LRT) to explore the genes with significantly differential expression levels across a series of time points (P < 10^−10^). (A) The stacked plot shows the number of upregulated (red) and downregulated (blue) DEGs of PAMs after being infected by SY18 (left) and HuB20 (right) strains, respectively. (B) The Venn diagrams show the shared genes in the two strains for upregulated (left) and downregulated (right) DEGs. (C) The heatmap of the DEGs with hierarchical clustering shows the expression levels of upregulated (red) and downregulated (blue) DEGs of PAMs infected by SY18 and HuB20 strains separately. The line plots illustrate the average trend of gene expression in hierarchical clusters.

To further investigate the expressional programs in the host response resulting from SY18 and HuB20 infection, we grouped the DEGs into a total of 28 clusters (13 clusters in SY18 strains and 15 clusters in HuB20 strains) of coexpressed genes based on their expression patterns (Fig 3C, S2 Table). A large fraction of DEGs in the upand downregulated groups, categorized as cluster Ⅰ, demonstrated linear expression changes along with time, implying cumulative effects of expressional changes in the host cell after virus infection. Additionally, we observed varying patterns of coexpressed gene clusters across different timepoints, suggesting that multiple dynamic transcriptional programs were involved in the host response.

Next we sought to identify the transcriptional program regulators, i.e., transcription factors (TFs) enriched in different clusters of coexpressed genes, through MEME motif search on the promoters of the selected genes (S3 Table). A number of TFs with known regulatory functions in the immune response, cytokines release, and type I IFN activation were identified, such as SP1, PATZ1, ETV5, STAT2, and IRFs (Fig S2). Interestingly, while some TFs, e.g., SP1, ETV5, STATs and IRFs, were enriched in multiple DEG clusters, other TFs showed enrichment in specific clusters. For example, ZN341, a transcriptional activator of STAT1 and STAT3 transcription, whose function was involved in the regulation of immune homeostasis, was enriched only in cluster II of SY18 host response genes. Our analysis demonstrates an intricate regulatory network for dynamic host response transcriptional programs.

### Pathway enrichment analysis of host DEGs reveals proinflammatory response after ASFV infection

To understand the pathways and biological processes enriched in the host transcriptome response to ASFV virus infection, we performed Gene Ontology (GO) enrichment analysis for the up(Fig 4A, S4 Table) and downregulated (Fig 4B, S4 Table) DEGs of the two virus strains respectively. As expected, in the upregulated group, DEGs of both strains were significantly enriched in immune and inflammation-associated pathways, including toll-like receptor (TLR) pathway, NF-kappaB transcription activity, cytokines production and interferon gamma production. Interestingly, when we look into the relationship between the genes and top upregulated pathways, we identified TLR genes such as TLR1, TLR3, and TLR7 as connector genes amongst different pathways (Fig 4B), highlighting the upregulation of TLRs as key genes in the host response to ASFV infection. Moreover, we noticed that in addition to TLR9 signaling which primarily recognizes DNA, TLR3 signaling, and which is located on the endosome membrane and primarily recognizes dsRNA, was also enriched. This is consistent with previous studies showing ASFV replication in the cytoplasm instead of the nucleus, a unique feature of ASFV compared to other DNA viruses (6, 7, 35). Thus, ASFV replication in the cytoplasm may be responsible for the activation of the TLR3 signaling, which might be a crucial step in inducing the innate immune response and inflammatory responses in the host.

**Fig 4.**
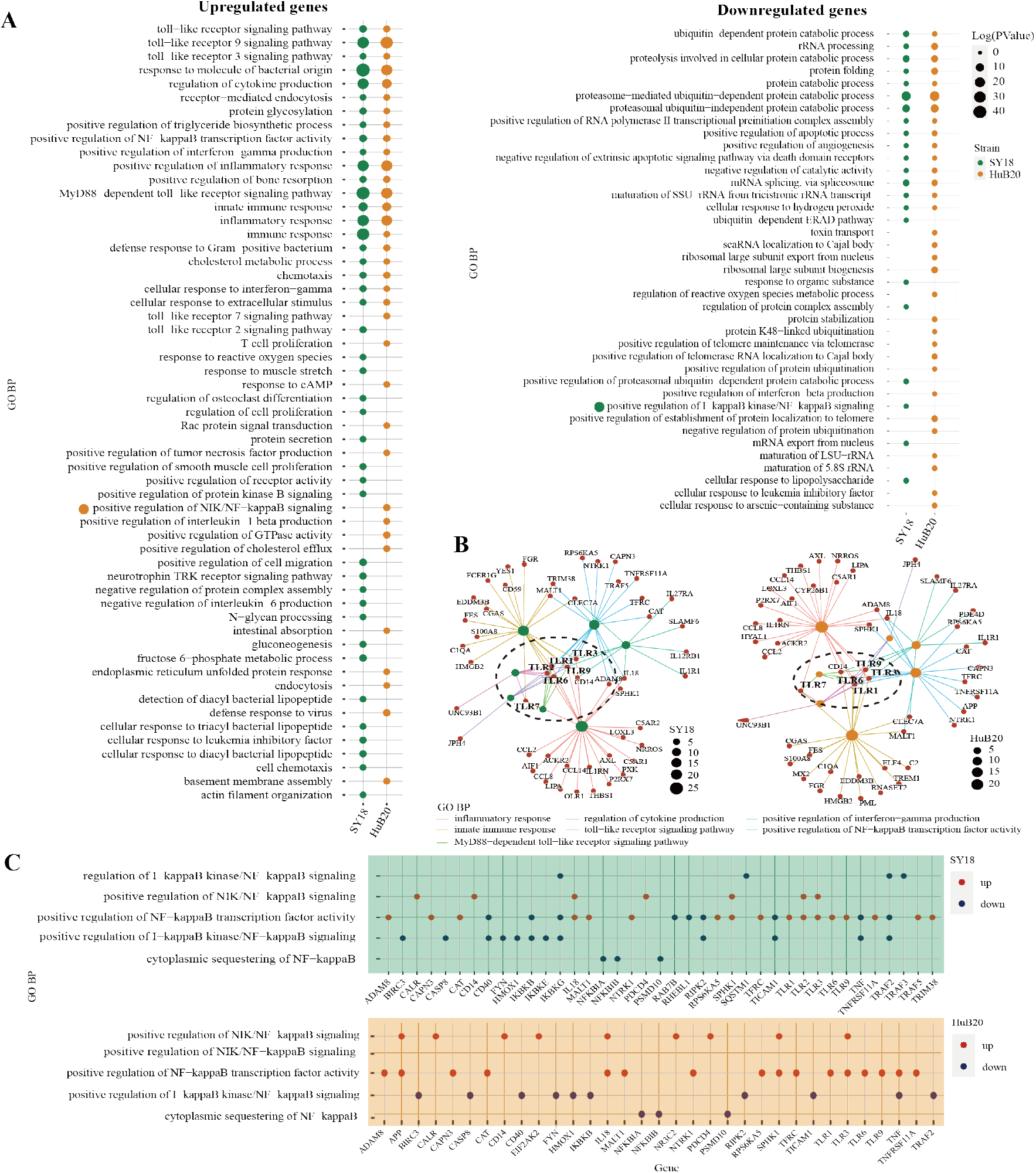
GO analysis of the genes with expression changes at 6, 12, 24, and 48 hpi. (A) Gene ontology biological processes (GO-BP) enrichment analysis of upregulated (left) and downregulated (right) DEGs (P < 10^−10^) of the two strains separately, and the bubble plot shows the GO terms with P < 0.01. (B) The network shows the relationship of most enriched up-regulated GO terms in PAMs after being infected by SY18 (left) and HuB20 (right) strains. (C) Dot plots of NF-kappaB related GO terms enriched by DEGs of PAMs after being infected by SY18 (top) or HuB20 (bottom) strains.

Meanwhile, DEGs in the downregulated panel in response to both virus strains were mainly involved in the proteasome-mediated protein catabolic process and apoptotic process, suggesting both ASFV strains were able to inhibit degradation of protein catabolic process and cell death through transcription. Notably, T cell proliferation, defense response to virus, response to cAMP, and positive regulation of interleukin-1 beta production were specifically enriched in the upregulated DEGs of HuB20 infected cells, whereas cell chemotaxis, protein secretion, and negative regulation of interleukin-6 production were enriched only in the upregulated DEGs of SY18 infected cells, indicating the two virus strains might stimulate different cytokines/chemokines response in the host.

To take a deeper dive into the gene-pathway relationships, we next plotted the involvement of all the DEGs related to NF-kappaB signaling in SY18 and HuB20, respectively (Fig 4C). NF-kappaB is known as a central pathway in the host cell in response to ASFV infection (36-38). While previous studies reported expression or activity changes in genes related to NF-kappaB, neither a clear picture of the NF-kappaB response to ASFV infection, nor the similarities and differences between strains of different virulence have been described. In our analysis, both SY18 and HuB20 enriched the same NF-kappaB related pathways and regulated gene expression changes in mostly the same directions (S5 Table). However, the exact DEGs involved, and the directional expression changes of DEGs were quite different. Our analysis reveals, for the first time, the differential activation of the essential NF-kappaB signaling between SY18 and HuB20 through differential expression regulation of NF-kappaB pathway genes.

### Diverse cytokines and chemokines responses induced by ASFV infection

Cytokines and chemokines-mediated immune and inflammatory responses are critical for ASFV pathogenicity (39, 40). Despite extensive efforts to study the differences in cytokines and chemokines expression after ASFV infection, results reported thus far remain contradictory (31, 32). To better understand the regulation of cytokines and chemokines by the two ASFV strains over time, we plotted the relative expression profiles of cytokines and chemokines DEGs, and validated their expression by RT-qPCR (Fig 5A, B). The cytokines and chemokines DEGs were grouped into three clusters with distinct patterns in expression. The first cluster contains mainly downregulated factors in both SY18 and HuB20 infected cells that are all involved in immune and inflammatory responses. We validated this finding through RT-qPCR, confirming that the expression of the proinflammatory factors IL-1β, CCL4, TNF and CXCL8 decreased progressively with viral infection (Fig 5B).

**Fig 5.**
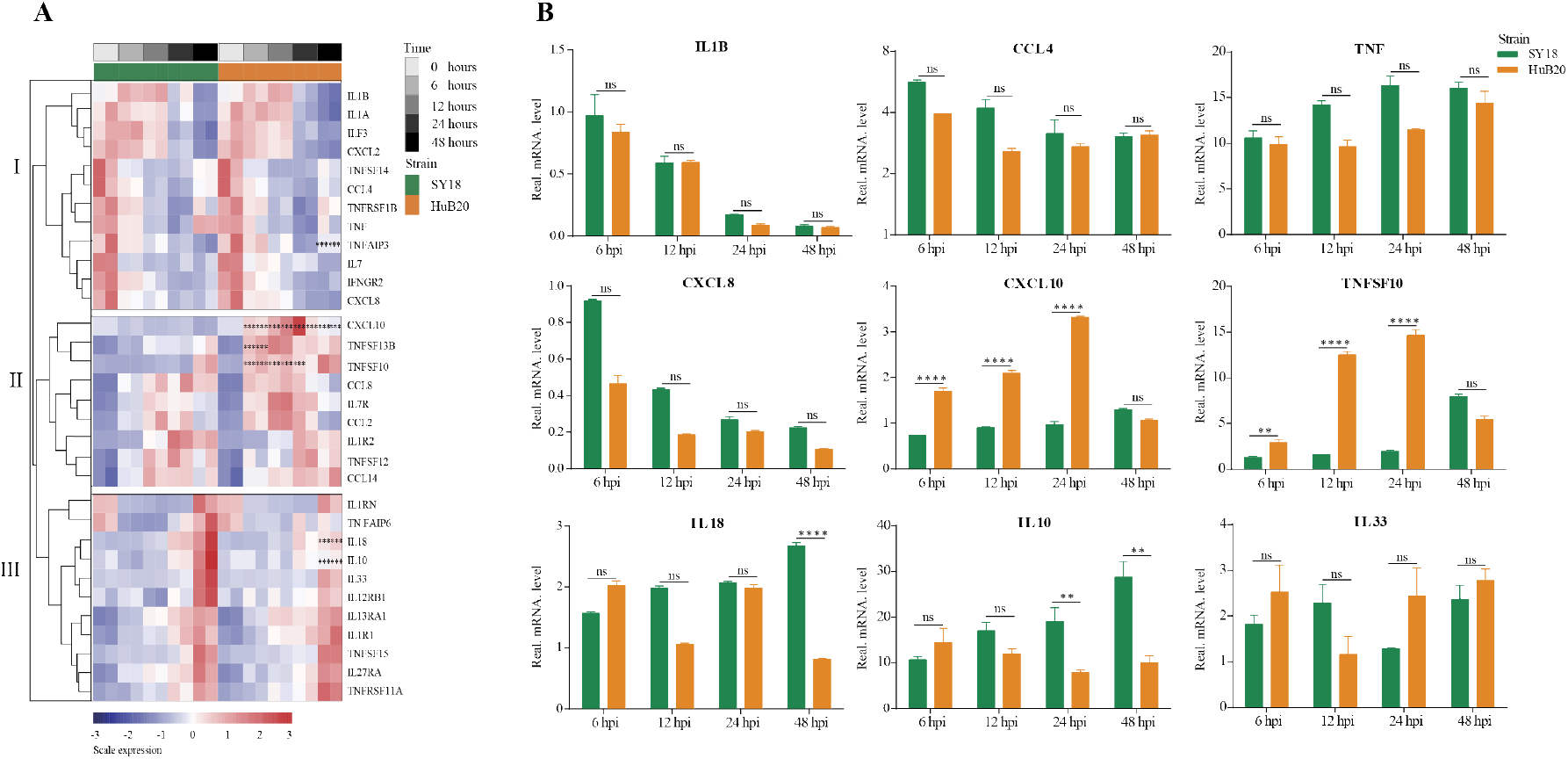
Patterns of cytokines changes and chemokines expression in PAM at different times after ASFV infection. (A) Heatmap of cytokines and chemokines expression after ASFV infection. cytokines and chemokines were divided into 3 clusters according to distinct patterns in expression over time. Dots illustrate the significance compared between two strains at the same time point with constraints the absolute value of log_2_foldchange > 1 and P < 0.1. *, P < 0.1; **, P < 0.05; ***, P < 0.01, ns, not significant. (B) Validation of randomly selected host cytokines and chemokines expression by real time-PCR at 6, 12, 24, and 48 h after PAMs infected by two ASFV strains (MOI= 3). Data are presented as mean ± SD of three independent experiments. The fold-difference was measured by the 2^-ΔΔCt^ method. Differences were assessed using a two-sample t-test. Significance was defined at P *<* 0.05 and log_2_foldchange >1. *, P < 0.05; **, P < 0.01; ***, P < 0.001, ****, P < 0.0001, ns, not significant.

The second and third cluster cytokines and chemokines genes were all upregulated after virus infection, including interleukins, interleukin receptors, TNF superfamily genes, and C-C motif chemokines. Note that several genes showed significant expression differences between SY18 and HuB20. In particular, CXCL10, TNFSF10, and TNFSF13B, critical regulator or effector genes in immune and inflammatory responses, showed significantly increased expression in HuB20 infection relative to SY18 from 6 hpi, suggesting that these genes might be responsible for the rapid induction of a stronger immune or inflammatory response in attenuated ASFV infection. On the contrary, increased expression of the inflammatory genes IL10 and IL18 at 48 hpi were significantly higher in SY18 compared to HuB20. These cytokines might contribute to the more severe tissue damage caused by the highly virulent strains in later stages of infection. Taken together, our cytokines and chemokines analysis revealed integrated and complex regulation of immune and inflammatory responses following ASFV infection. Our analysis suggests differential expression of cytokines and chemokines factors, such as IL10, IL18, CXCL10 and TNFSF10, may be associated with the differential pathogenicity of the two ASFV strains with different virulence.

### Stronger innate immune response stimulated by HuB20 than SY18

To further explore the differential expression program in the host response between SY18 and HuB20 along the infection timeline, we considered the interaction term of virus strains and time points and fitted a Likelihood Ratio Test to identify differentially expressed genes. A total of 6 clusters with at least 15 genes with similar expression patterns were found (Fig 6A, S6 Table). GO enrichment analysis of individual clusters identified cluster Ⅰ and Ⅱ were enriched in innate immune response related biological processes such as type I interferon signaling, interferon-stimulated gene 15 (ISG15)-protein conjugation and the JAK-STAT cascade (S7 Table). Interestingly, starting from 6 hpi, DEGs in cluster Ⅰ and Ⅱ were rapidly upregulated in HuB20 infected cells while the expression of these innate immune response related genes changed gradually in SY18 infected cells (Fig 6B), suggesting that the low virulent HuB20 strain could stimulate a rapid immune and defense response in the early stages of infection. In addition, we observed high transcript levels of intracellular sensors and receptors (cGAS, P < 3.82e^-12^, PARP9, P < 1.35e^-2^, CD38, P < 7.85e^-5^ IFIH1/MDA5, P < 2.05e^-15^ and FCGR1A, P < 9.96e^-26^) from 6 hpi of HuB20, whose functions were related to the recognition of viral DNA as well as viral RNA (41-45).

**Fig 6.**
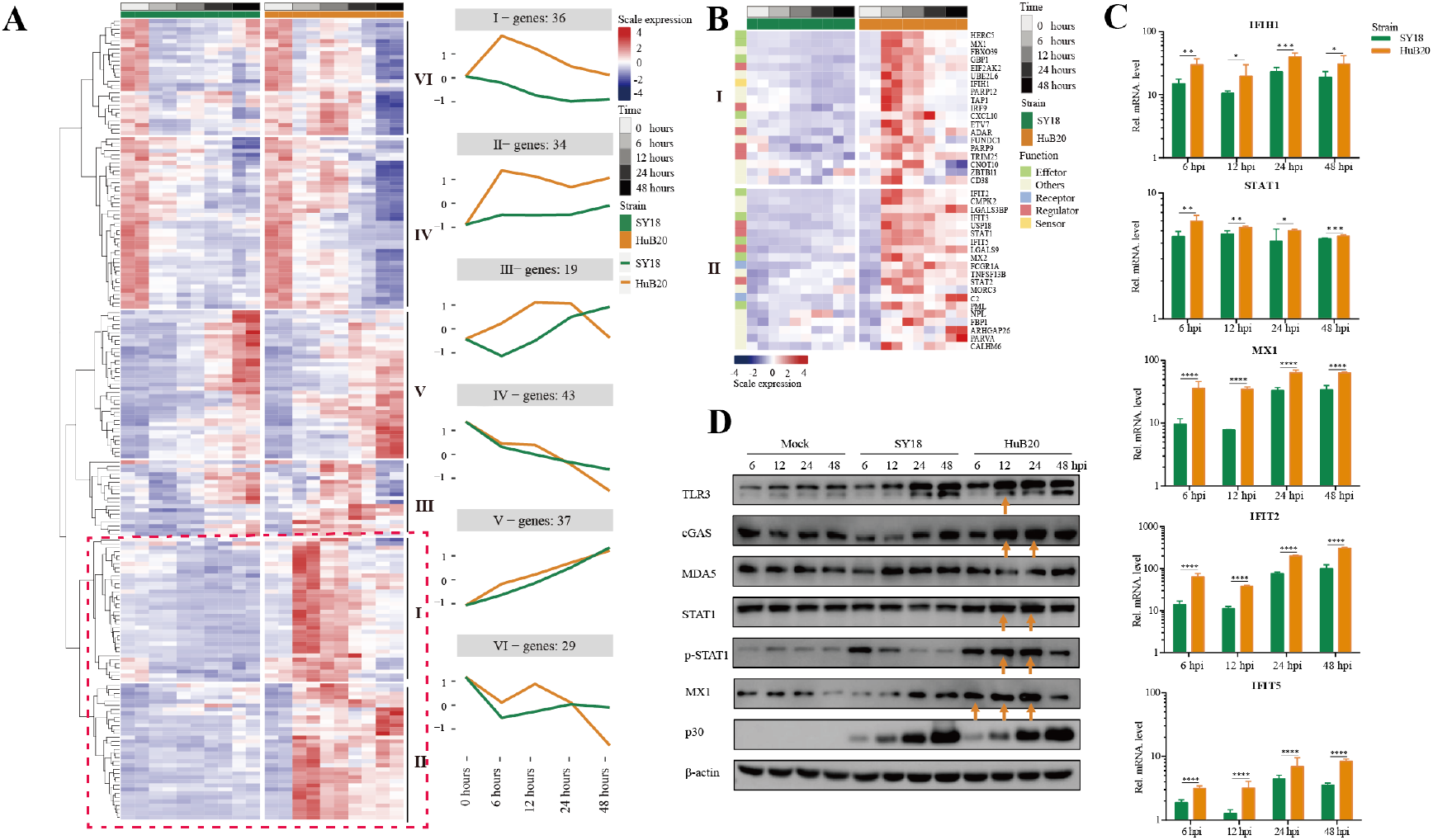
Comparison of host gene expression differences. (A) Heatmap of DEGs considering the effects of infection of SY18 and HuB20 separately over time on PAMs with the LRT test. The full model’s design formula includes the effects of infection over time, and the reduced model removes this term to perform the LRT test. The line plots illustrate the average trend of gene expression in clusters. Each cluster has at least 15 genes. (B). Heatmap of the DEGs in clusters I and II, with immune related functions annotated for each gene. (C) Validation of innate immunity associated gene expression by real time-PCR. PAMs were infected or mocked infected by ASFV SY18 and HuB20 strains, respectively (MOI= 3), at 6, 12, 24 and 48 hpi. Total RNA was extracted from the PAMs and subjected to RT-qPCR to quantitate IFIH1, STAT1, MX1, IFIT2 and IFIT5 expression. The data were normalized using β-actin. The fold-difference was measured by the 2^-ΔΔCt^ method. Differences were assessed using a two-sample t-test. Significance was defined at P <0.05. *, P < 0.05, **, P < 0.01; ***, P < 0.001, ****, P < 0.0001, ns, not significant. (D) Western blotting analysis of innate immunity associated proteins. PAMs were infected or mocked infected by ASFV SY18 and HuB20, respectively (MOI= 3), at 6, 12, 24 and 48 hpi. Cell lysates were collected and subjected to Western blotting analysis using the indicated antibodies.

Furthermore, we noted that STAT1, STAT2, IRF9, USP18 and TRIM25 also exhibited high levels of transcription earlier in HuB20 infection than SY18. Phosphorylated STAT1 and STAT2, together with IRF9 are known to form the interferon-stimulated gene factor 3 (ISGF3) complex, which transcriptionally activates the ISGs (46, 47). In addition, the RT-qPCR results also proved that HuB20 infection induced higher levels of innate immune-related factors in PAMs than SY18 infection (Fig 6C). Therefore, we suspect that activation of these regulators might contribute to the introduction of a rapid immune response by the attenuated ASFV strain. We further validated the consistency of transcriptome changes relative to the protein level using Western blotting (Fig 6D). Albeit delayed activation at protein level compared to transcript level, the innate immune response genes including cGAS, STAT1/pSTAT1 and MX1 all demonstrated high levels of activation in HuB20 infected cells compared to SY18 (Fig 6D). Taken together, we identified a subset of genes that were activated rapidly after infection with the low virulent HuB20 strain, and demonstrated for the first time, that the innate immune response involving cGAS pathway, JAK-STAT and IFN stimulated genes in the host cells were activated quickly after infection with the low virulent strain HuB20, while these pathways were immune escaped by the highly virulent strain SY18.

### Host-virus coexpression network reveals new insights into the functions of viral genes

To explore the relationship between host and virus gene expression dynamics, we constructed a host-virus coexpression network for each of the ASFV strains (Fig 7, S8 Table). First of all, we observed a module in the SY18 network with multiple viral genes sharing similar connection to a set of host genes (Fig 7A). In particular, I196L as a hub gene shared 88% (15/17) of host coexpression with NP868R, suggesting these two viral genes might be coregulated or involved in similar interactive processes with host cells.

**Fig 7.**
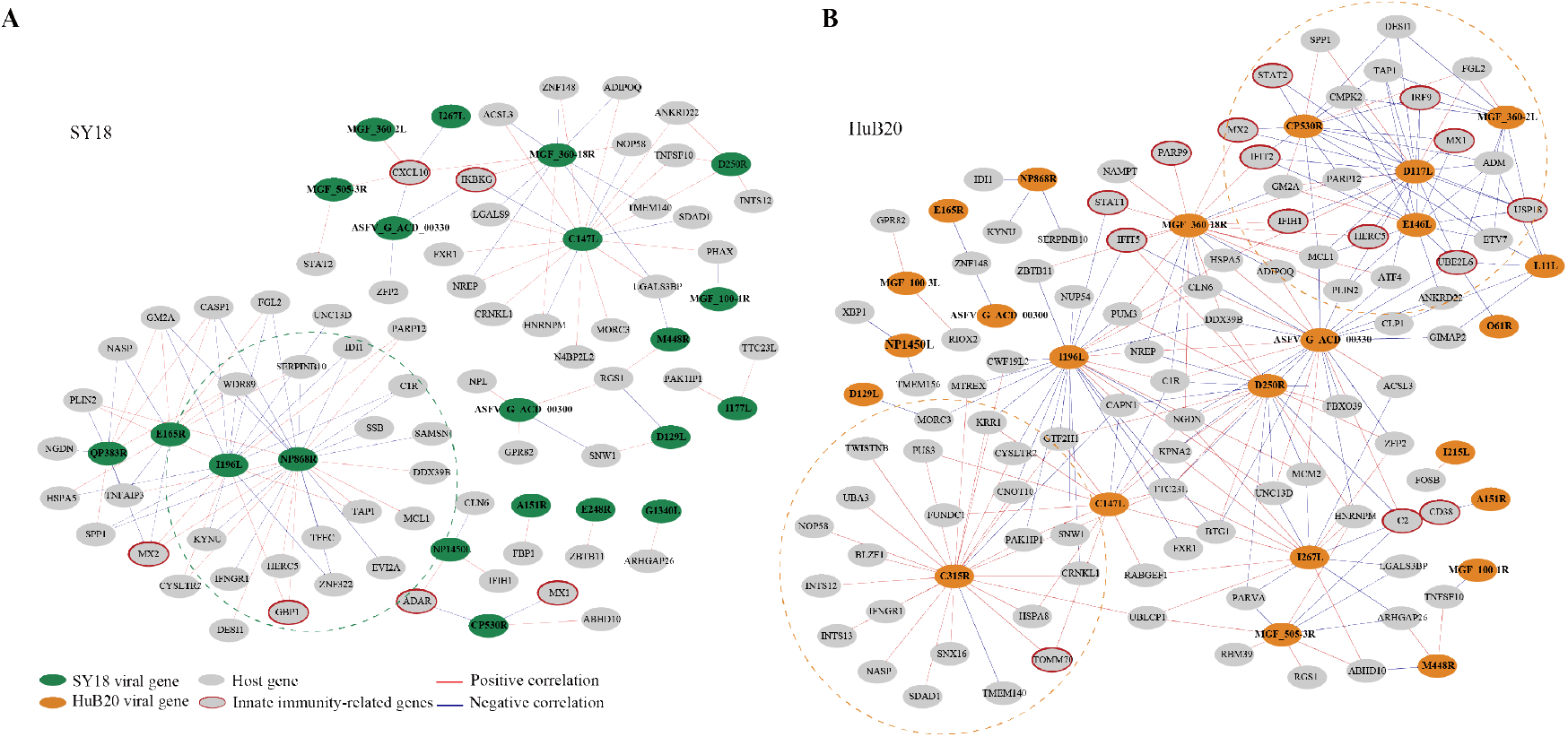
Correlation between ASFV and host genes. The correlation between the ASFV and host DEGs was measured using Pearson correlation coefficients for the corresponding gene expression and is visualized in the network for SY18 (left) and HuB20 (right) strains. The absolute value of the Pearson correlation coefficient > 0.9 was considered significant. Red and blue lines indicate positive and negative correlations, respectively.

Secondly, we noticed that in the HuB20 network (Fig 7B), a module involving MGF_360-2L, CP530R, E146L and D117 viral genes was negatively correlated with innate immunity-related genes, e.g. IRF9, USP18, UBE2L6, IRF9, STAT2, MX1/2, IFIT2 and HERC5. Among these viral genes, MGF_360-2L has been shown to be involved in the pathogenicity of ASFV in pigs, where deletion of multiple MGF360 family genes increased the expression of ISG and type I IFNs in infected macrophages (23, 24, 48). However, CP530R, E146L and D117 have never been reported to be associated with innate immunity, but the expression levels of these genes in HuB20 infected cells was all significantly lower compared to SY18 (Fig S3), indicating lower expression of these genes might account for the activated immune response in HuB20 infected cells.

In addition, the viral gene C315R, which encodes TFIIB-like transcription factor, is involved in the regulation of RNA transcription and modification (49). Indeed, we identified positive correlation of C315R with RNA polymerase I subunit F (TWISTNB), integrator complex subunit 12 (INTS12) and mtr4 exosome RNA helicase (MTREX) transcription, which were involved in RNA processing and splicing. Intriguingly, C315R was also associated with genes involved in protein transport between the endoplasmic reticulum (ER) and Golgi, such as NOP58 ribonucleoprotein (NOP58), basic leucine zipper nuclear factor 1 (BLZF1), sorting nexin 16 (SNX16), SDA1 domain containing 1 (SDAD1) and nuclear autoantigenic sperm protein (NASP), indicating possible functional association of C315R with protein transport of the host cells.

Lastly, the same viral genes (e.g. MGF_360-18R, I196L and C147L) often associate with different host genes between SY18 and HuB20 infected cells, suggesting a highly dynamic interactive relationship between the virus and host expression programs. Therefore, elaborating the transcriptional correlation between host and virus genes might provide novel insights to explore the unknown functions of some viral genes, or provide a reference map for target selection to guide vaccine or drug development for ASF disease.

## Discussion

RNA sequencing (RNA-seq) has been applied to study various biological processes, such as revealing the interaction of virus infection and host response (10, 13, 50). However, studies using RNAseq to preform transcriptomic profiling of ASFV and infected host cells are scarce, and these studies target a single strain or time point and do not provide a comprehensive picture of host-virus interactions (8, 14-17). Here, we integrated RNA-seq analysis to examine and compare the transcriptomic landscape of porcine PAMs during infection with highly (SY18) and low virulent (HuB20) ASFV strains at different stages of infection, depicting unprecedented details about the temporal host response after ASFV infection. By combining functional enrichment analysis and experimental validation, we highlight similarities and differences in viral and host gene expression patterns and cellular pathways. In particular, we elucidated differences in host innate immune and inflammatory responses stimulated by ASFV, which may provide novel insights for intensive study of ASFV pathogenicity and therapeutic targets.

Several transcriptomic and other experimental studies have shown that ASFV infection leads to changes in the transcription of pattern recognition receptors (PRRs) in some TLR signaling pathways, as well as significant changes in the transcription of some antiviral and inflammatory factors (21, 26, 51, 52). Our data show that the upregulation of multiple TLRs (e.g. TLR1, 3, 7) acts as connectors mediating the regulation of multiple responses, especially cytokines and chemokines production and innate immune signaling (Fig 4C). Furthermore, published literature has shown that infection with ASFV of different virulence can lead to differential inflammatory responses, immune responses and apoptotic processes, while the relevant mechanisms remain unclear. Interestingly, in our results, we noticed that DEGs in both SY18 and HuB20 infection were enriched to the NF-kappaB signaling pathway. By comparing the unique DEGs involved in NIK/NF-kappaB signaling and predicting their enrichment in other known pathways (Fig 4C, S6 Table), we found substantial differences between SY18 and HuB20, which may account for the different host responses they elicited. The above results provide new insights and research targets into the role of NF-kappaB-regulated immune, inflammatory and apoptotic responses in ASFV infection. Certainly, further experimental data to confirm these observed relationships will facilitate the study of the mechanisms by which ASFV regulates host responses

We also analyzed the expression patterns of cytokines and chemokines and performed experimental validation by RT-qPCR after ASFV infection, revealing differences in the expression patterns of related factors caused by different ASFV strains, providing a theoretical basis for the study of ASFV pathogenicity. By comparing the differential expression programs of SY18 and HuB20 in the host response, we found that HuB20 provokes stronger host immune response at early stage than SY18, which was supported by the quickly activated high expression of receptor, sensor and regulator genes. In particular, the correlation network between the viral and host gene expression suggests a clear relationship between the HuB20 viral genes (e.g. CP530R, D117L, E146L and MGF_360-2L) and innate immunity genes. Our analysis demonstrates that deciphering the relationship between virus and host genes would contribute to resolving the unknown functions of ASFV genes and deepen the investigation of host-virus interactions.

Our study was limited by a single viral infection dose, within sample cell heterogeneity, individual gene variability and other confounding factors, such as annotation of the reference genome. However, we compensated for the differences caused by individual cell heterogeneity to some extent by comparing and analyzing the gene expression patterns of the two virulent strains over time as well as the overall regulatory pathway changes. Meanwhile, combined with previous studies, we analyzed and presented predictive results for a comprehensive set of regulatory pathways and persuasive targets of action following ASFV infection, which will provide insightful information for further investigations to understand the host response after ASFV infection and valid information for screening candidate targets for ASFV inhibition. Future ASFV related genomic datasets could provide the research community with important resources for ASFV studies.

## Materials and methods

### Cells, viruses and antibodies

Porcine alveolar macrophages (PAMs) were prepared from 2-month-old piglets bronchoalveolar lavage as described previously, cultured in Roswell Park Memorial Institute (RPMI) 1640 medium (Gibco, USA), supplemented with 10% fetal bovine serum (Gibco, USA) and grown at 37°C under 5% CO_2_ atmosphere. The ASFV SY18 strain (GenBank accession no.MH766894), a field virulent ASFV, was originally isolated from specimens in the initial ASF outbreak in China (53). The ASFV HuB20 strain (GenBank accession no.MW521382), a natural attenuated ASFV was isolated from the tissues of pigs in Hubei, China (34). The two viruses were passaged in primary PAMs and maintained at -80°C in the biosecurity level 3 lab. The monoclonal antibodies for cGAS, TLR3, STAT1 and p-STAT1 were purchased from Santa Cruz Biotechnology, USA, and anti-β-actin, IFIH1/MDA5 and MX1 were purchased from Proteintech Biotechnology, USA.

### Sample Preparation for RNA-sequencing

PAMs (10^6^ per well) were seeded in 6-well plates and mock-infected or infected with indicated ASFV strains at a multiplicity of infection of 3. After 1 hour of incubation, the viruses were removed, the cells were washed twice with PBS, and fresh 1640 medium was added. At the specified time points (0, 6, 12, 24 and 48 hpi), cells were harvested for RNA extraction

### Library preparation and RNA sequencing

The cDNA libraries were prepared according to the standard Illumina protocol (NEBNext® Ultra™ II RNA Library Prep Kit for Illumina®). Briefly, total RNA from the specified PAMs was treated with RNase-free DNase I (Vazyme, China) following the manufacturer’s instructions. The total amount and integrity of RNA was estimated using the Bioanalyzer 2100 system (Agilent Technologies, USA) with the RNA Nano 6000 Assay Kit. First strand cDNA was synthesized using random hexamer primers and M-MuLV Reverse Transcriptase, and then RNaseH was used to degrade the RNA. Second strand cDNA synthesis was subsequently performed using DNA Polymerase I and dNTPs. After commencing PCR amplification, the PCR product was purified by AMPure XP beads, and the library was finally obtained. The libraries were quantified by an Agilent 2100 bioanalyzer and then subjected to sequencing using an Illumina NovaSeq 6000 sequencer.

### Cell total RNA isolation and real-time quantitative PCR (RT-qPCR)

PAMs in 6-well plates were infected with ASFV SY18 and HuB20 at 3 MOI (multiplicity of infection), respectively. Mock-infected cells were used as control. Cells were collected at 6, 12, 24 and 48 hours post inoculation, and total RNA was isolated using a Total RNA Kit I (Omega Bio-Tek, USA) and reverse transcribed with HiScript Ⅱ Q RT SuperMix (Vazyme, China) according to the manufacturer’s instructions. qPCR was performed using ChamQ Universal SYBR qPCR Master Mix (Vazyme, China) on a LightCycler® 480 Real-Time PCR System. The relative expression of mRNA was calculated based on the comparative cycle threshold (2^^-ΔΔCt^) method and normalized with porcine β-actin mRNA levels. The primer sequence information is provided in S9 Table. The results were analyzed using GraphPad Prism 6 software.

### Western blotting analysis

Cell lysates were subjected to 10% SDS-PAGE and then transferred to nitrocellulose (NC) membranes. The membranes were blocked for 1.5 h at room temperature in Tris-buffered saline containing 10% nonfat dry milk and 0.05% Tween 20 (1×TBST) and were incubated at 4°C for 12 h with the indicated antibodies. The Membranes were washed with 1×TBST, incubated with horseradish peroxidase (HRP)-labeled goat anti-rabbit IgG or anti-mouse IgG antibody (Beyotime, China) at room temperature for 45 min, and treated with enhanced chemiluminescence (ECL) reagent (Thermo Fisher Scientific, USA).

### RNA-seq data quality control and read mapping

Raw data (sequencing reads) in fastq format were first processed through in-house Perl scripts. In this step, clean data (clean reads) were obtained by removing reads containing adapters, reads containing N bases and low-quality reads from the raw data. All downstream analyses were based on cleaned data (v0.20.1) (54). The clean reads were aligned to the Sus Scrofa Largewhite_V1 (GCA_001700135.1) and ASFV (SY18 (GenBank accession no.MH766894) and HuB20 (GenBank accession no.MW521382)) genome assemblies using STAR (v2.7.8a) with default parameters (55). Uniquely mapped read pairs were counted using featureCounts (v2.0.1) (56). To make the count matrix more comparable among samples, normalization is the critical process of scaling raw counts. Hence, the count matrix was normalized based on the median of ratios method using the R package DESeq2 (v1.32.0), and the rlog transformation was performed for PCA plots (57).

### Differential expression analysis

Differential expression analysis was performed using DESeq2 exploiting the likelihood ratio test (LRT) and testing a full and reduced formula for time-course analyses for each strain of the virus separately with a P < 10^−10^ (Fig 3). LRT is used to explore whether there are significant differences in the effect of the timeline. However, for those differentially expression genes, the expression variations were not consistent. Hence, after filtering genes that were significantly different over time, we clustered the genes using DEGreport (v1.28.0) in R to sort genes into groups based on shared expression patterns (60). In addition, we compared the gene expression levels of PAM cells infected by two strains at each time point to explore significant genes with padj < 0.1 and the absolute value of log_2_FoldChange >= 1 by DESeq2 Wald Test (Fig 5).

We are also interested in the differences in gene expression between SY18 and HuB20 infection over time. In other words, we want to compare expression levels by considering two conditions, virus and time, at the same time. Hence, we use a design formula for our full model to explore the difference between the two strains in the effect of the infection over time (virus: time). To perform the LRT test as we used before, we also need a reduced model without the interaction term virus: time. After applying the LRT test, significant genes were identified using a threshold padj < 0.01 (Fig 6). Differentially expressed viral genes were identified using a similar method, and the Pearson correlation coefficient between the host significant gene and the viral significant gene was computed.

Further analysis of ASFV genes used their noted or predicted functions, from the VOCS tool database (https://4virology.net/) entries for the SY18 and HuB20 genomes.

### Enrichment analysis of differentially expressed genes

Gene ontology (GO) analysis was performed on the differentially expressed genes on the DAVID 2021 website with default parameters using direct GO biological process categorization (58). Then we displayed the most significantly enriched terms (P < 0.01) in bubble plots. However, to better understand the potential biological complexities in which one gene may involve multiple categories, we used the tool provided by clusterProfiler (v4.0.5) in R to depict the linkage of genes and GO terms as a network (59).

### Promoter Motif Analysis

Two hundred bases upstream and fifty bases downstream of the transcriptional start site (TSS) sequences of genes in each cluster (13 clusters in SY18 and 15 clusters in HuB20) were extracted from the large white genome (GCA_001700135.1) in FASTA format using BEDtools. Sequences were analyzed in MEME (http://meme-suite.org) by using Human DNA motif database (HOCOMOCOv11 core HUMAN mono meme format) for enrichment. The output from MEME is a list of the sequences for which the E-value is less than 10. For the site positional distribution diagrams, all sequences are aligned on their centers. Position frequency matrices (PFMs) of motifs of interest were drawn by the R package motifStack (v1.40.0) (60).

## Supporting information

**S1 Table. ASFV gene counts and functional annotation. (XLSX)**

**S2 Table. Host gene counts and DEGs with time series. (XLSX)**

**S3 Table. Host gene promoter motif. (XLSX)**

**S4 Table. GO BP enrichment of host DEGs with time series. (XLSX)**

**S5 Table. Host DEGs involved in NF-kB related pathways. (XLSX)**

**S6 Table. Classification of DEGs between two strains across time points. (XLSX)**

**S7 Table. GO BP enrichment of DEGs in clusters Ⅰ and Ⅱ between the two strains across time points. (XLSX)**

**S8 Table. Pearson correlation between viral DEGs and host DEGs. (XLSX)**

**S9 Table. Primers for real time-PCR validation**.

## Acknowledgments

Thanks for the technical support by the core facilities of Zhejiang University Medical Center and Liangzhu Laboratory.

## Author Contributions

N.S. conceived the study; N.S., Z.S., T.C., and L.L. designed the experiment. L.L. and Y.Z. performed the experiments. T.Z., A.A., and X.Z. analyzed the data. N.S., Z.S., and T.C. directed the study. N.S., L.L., and T.Z. wrote the paper.

## Financial Disclosure Statement

This work is funded by the Starting Fund from Zhejiang University and Shandong Major Project of the Science and Technology Innovation (2019JZZY020606).

## Notes

### Competing Interest Statement

The authors have declared no competing interest.

